# SARS-CoV-2 Infection Impacts Carbon Metabolism and Depends on Glutamine for Replication in Syrian Hamster Astrocytes

**DOI:** 10.1101/2021.10.23.465567

**Authors:** Lilian Gomes de Oliveira, Yan de Souza Angelo, Pedro Yamamoto, Victor Corasolla Carregari, Fernanda Crunfi, Guilherme Reis-de-Oliveira, Lícia Costa, Érica Almeida Duque, Nilton Barreto dos Santos, Glaucia Maria Almeida, Egidi Mayara Firmino, Isadora Marques Paiva, Carolina Manganeli Polonio, Nagela Ghabdan Zanluqui, Marília Garcia de Oliveira, Gustavo Gastão Davanzo, Marina Caçador Ayupe, Caio Loureiro Salgado, Antônio Francisco de Souza Filho, Marcelo Valdemir de Araújo, Taiana Tainá Silva-Pereira, Angélica Cristine de Almeida Campos, Luiz Gustavo Bentim Góes, Marielton dos Passos Cunha, Maria Regina D’Império Lima, Denise Morais Fonseca, Ana Márcia de Sá Guimarães, Paola Camargo Minoprio, Carolina Demarchi Munhoz, Cláudia Madalena Cabrera Mori, Pedro Manoel Moraes-Vieira, Thiago Mattar Cunha, Daniel Martins-de-Souza, Jean Pierre Schatzmann Peron

## Abstract

Coronaviruses belong to a well-known family of enveloped RNA viruses and are the causative agent of the common cold. Although the seasonal coronaviruses do not pose a threat to human life, three members of this family, i.e., SARS-CoV, MERS-CoV and recently, SARS-CoV2, may cause severe acute respiratory syndrome and lead to death. Unfortunately, COVID-19 has already caused more than 4.4 million deaths worldwide. Although much is better understood about the immunopathogenesis of the lung disease, important information about systemic disease is still missing, mainly concerning neurological parameters. In this context, we sought to evaluate immunometabolic changes using *in vitro* and *in vivo* models of hamsters infected with SARS-CoV-2. Here we show that, besides infecting hamster’s astrocytes, SARS-CoV-2 induces changes in protein expression and metabolic pathways involved in carbon metabolism, glycolysis, mitochondrial respiration, and synaptic transmission. Interestingly, many of the differentially expressed proteins are concurrent with proteins that correlate with neurological diseases, such as Parkinsons’s disease, multiple sclerosis, amyotrophic lateral sclerosis, and Huntington’s disease. Metabolic analysis by high resolution real-time respirometry evidenced hyperactivation of glycolysis and mitochondrial respiration. Further metabolomics analysis confirmed the consumption of many metabolites, including glucose, pyruvate, glutamine, and alpha ketoglutarate. Interestingly, we observed that glutamine was significantly reduced in infected cultures, and the blockade of mitochondrial glutaminolysis significantly reduced viral replication and pro-inflammatory response. SARS-CoV-2 was confirmed *in vivo* as hippocampus, cortex, and olfactory bulb of intranasally infected hamsters were positive for viral genome several days post-infection. Altogether, our data reveals important changes in overall protein expression, mostly of those related to carbon metabolism and energy generation, causing an imbalance in important metabolic molecules and neurotransmitters. This may suggest that some of the neurological features observed during COVID-19, as memory and cognitive impairment, may rely on altered energetic profile of brain cells, as well as an unbalanced glutamine/glutamate levels, whose importance for adequate brain function is unquestionable.

## INTRODUCTION

Human coronaviruses are enveloped 30Kb single-stranded (+)RNA viruses and the causative agent of common cold and severe acute respiratory syndrome (SARS). Six members of the *coronaviridae* family are capable of infecting humans, considered of low (HCoV 229E, HCoV NL63, HCoV HKU1 and CoV OC43) or high (SARS-CoV, MERS-CoV and SARS-CoV2) pathogenicity^1^. SARS-CoV-2 pandemic started in December 2019 in Wuhan, China after the probable spill over from bats due to mutations of the spike proteins^2, 3^.

Mild disease is observed in most patients, who develop fever, headache, and coughing. Some patients, however, evolve to severe acute respiratory syndrome (SARS), characterized by lung inflammation with intense myeloid cells inflammatory infiltrate, hyaline membrane formation and alveolitis, clearly evidenced by lung magnetic resonance imaging (MRI)^2, 4, 5^. This results in dyspnea, difficult breathing, and low oxygen saturation. Most severe cases evolve with cytokine storm and systemic organ failure^6^, mainly in people with comorbidities such as hypertension, obesity, and diabetes, probably either to altered immune responses or to the higher expression of angiotensin converting enzyme-2 (ACE-2) in the lungs ^7^. Since January 30^th^, 2020, when the WHO declared a state of emergency and international concern, more than 5,7 million people have died of COVID-19.

Viral particles invade host cells using its Receptor Binding Domain (RBD) of the spike protein through the interacting with Angiotensin Converting Enzyme-2 (ACE-2), expressed mostly on type II pneumocytes but also on other cell types, including neurons, enterocytes and many others^8, 9^. Further, spike is cleaved in subunits S1 and S2 by the protease TMPRSS2^10^ on the surface of host cells, promoting the fusion with cell membrane for further release of the genome into the cytoplasm. In the cytoplasm, ORF1a and ORF1b are translated from the (+) RNA strand into 16 non-structural proteins, whereas (-) RNA strands are used as template for replication and for the translation of sub-genomic RNAs. Sub-genomic RNAs are then translated into structural proteins, as spike (S), envelope (E), nucleocapsid (N) and membrane (M), that are assembled into viral particles and released to the extracellular space^11^. During this process, viral RNA may be sensed by innate immunity sensors, as RIG-I^12^ and MDA-5^13, 14^ leading to inflammasome activation and cytokine secretion. It triggers signaling pathways that culminate in the production of pro-inflammatory and anti-viral cytokines, as IL-1b, IL-6, TNF-α and type I interferons, which are exacerbated in patients evolving to cytokine storm, as reviewed^15^. Conversely, this is associated with intense lung infiltration of myeloid cells, such as neutrophils^16^ and monocytes, as well as of lymphocytes with high expression of the exhaustion markers, PD-1 and TIM-3^17^. Clearly, the immune response during COVID-19 has a dual role, as it may reduce viral burden, but also account for disease exacerbation^18^. These evidences show the complexity of COVID-19 as well as the difficulty in predicting worse disease outcomes.

In this context, despite being a respiratory disease, COVID-19 patients may develop several other manifestations^18^, including anosmia and ageusia^19^, suggesting an impact in the central nervous system. In fact, it has been demonstrated that SARS-CoV-2 infects olfactory sensory neurons (OSN) and uses the transmucosal route to reach brain tissue^20^. The neurological importance of SARS-CoV-2 infection is still a matter of debate. A retrospective study performed in Wuhan, China, from January to February 2020 analyzed 214 patients, of whom 36.4% developed neurological symptoms related to cerebrovascular events and impaired consciousness^21^. Since then, many case reports and cohort prospective studies have been published pointing to signs and symptoms of neurological alterations that range from the anosmia and ageusia, to leptomeningitis^22^, stroke, microvascular damage^23^, severe headache, encephalopathy^24^, and even peripheral Guillain-Barré Syndrome^25^. Histopathological analysis revealed multiple ischemic areas, encephalopathy, and microglial activation. Interestingly, recent reports have also pointed to residual psychiatric features in patients that recovered from COVID- 19, as chronic fatigue, cognitive decline^26^, mood disorders and depression, in the so-called “brain fog” ^27, 28^.

Whether these are direct viral effects or indirect results of systemic inflammation is a matter of controversy, and the real impact of SARS-CoV-2 to the human brain remains elusive. In this sense, here we used *in vitro* and *in vivo* systems to evaluate immunometabolic and energetic changes using Golden Syrian Hamsters (*Mesocricetus auratus*) infected with SARS-CoV-2. We observed that the virus infects and replicates in hamster’s astrocytes and further inducing a pro-inflammatory response. Interestingly, we observed that SARS-CoV-2 shifts main metabolic pathways, specially subverting the glutamine usage for enhanced replication. Corroborating this, the blockade of glutaminolysis significantly impairs viral progeny. Also, respirometry confirmed intense metabolic activity in the presence of the virus, further corroborated by metabolomics. *In vivo*, we confirmed the presence of viral RNA in the cortex and hippocampus of infected animals, and consistent changes in the protein profile. Thus, here we suggest that the virus hijacks cellular carbon metabolism, especially glutamine consumption, for its own benefit, in detriment of a proper cellular metabolic function. Due to the importance of glutamatergic synapses for appropriate neuronal synapses, here we propose a correlation between SARS-CoV-2 infection over memory and cognitive impairment due to metabolic imbalance of important neurotransmitters, especially glutamate/glutamine. Moreover, our proteomic analysis showed a significant impact on carbon metabolism pathways consistent with brain diseases, such as Parkinson’s Disease, Multiple Sclerosis, and long-term depression, which were also concordant with changes observe in COVID-19 deceased patients. This brings attention to a possible correlation of altered carbon and energy metabolism with the cognitive and memory impairment observed in COVID-19 patients.

## MATERIALS AND METHODS

### Virus isolation and propagation

SARS-CoV-2/SPPU-0691 (GenBank accession number MW441768) was obtained, isolated, and propagated at Scientific-Platform Pasteur-USP (SPPU) and used for the experiments *in vitro*. The respiratory sample (naso-oropharyngeal swab) was acquired during the acute phase of infection and complying with the inclusion criteria for clinical suspicion of associated viral infection to SARS-CoV-2 adopted by the Ministry of Health of Brazil. In sum, viral stocks were generated in Vero CCL81 cells. Prior to infection, cells were maintained in DMEM Low glucose (LGC^®^) supplemented with 10% of fetal bovine serum (FBS) (Gibco^®^) and 1% penicillin/streptomycin (LGC^®^) at 37°C, 5% CO_2_. The initial inoculum (passage 1) was prepared to dilute the clinical samples (1:5) in non-supplemented DMEM low glucose (Gibco^®^). The inoculum was then added to the flask containing Vero cells and maintained for 1 hour at 37°C, following addition of fresh DMEM low glucose media supplemented with 2% FBS and 1% penicillin/streptomycin. The cell culture was observed for cytopathic effects on each day after the inoculation. When a prominent cytopathic effect was observed the cell culture supernatant was collected, centrifuged to remove debris (500g), and stored at -80°C as the first viral passage. The same strategy was employed three times to obtain a fourth passage isolate. Fourth passage viral stocks were used in this study titrated by plaque-forming unit (PFU) assay, according to protocols described elsewhere^23^. SARS-CoV-2 strain human/BRA/SP02cc/2020 (GenBank access number: MT350282.1) was used in the *in vivo* experiments.

### Primary astrocytes cultures of Golden Syrian Hamsters (*Mesocricetus auratus*)

Primary astrocytes were obtained from Syrian hamster neonates at days 2-4 postpartum (dpp). The protocols were approved by the Ethics Committee for Animal Research of University of Sao Paulo (CEUA n° 7971160320 / 3147240820) and all efforts were made to minimize animal suffering. Animals were obtained from the Department of Pathology, School of Veterinary Medicine and from the Animal Science Department of Preventive Veterinary Medicine and Animal Health from the University of São Paulo. Briefly, brains were harvested, and the meninges were removed following tissue digestion with 0.05% of trypsin (Gibco^®^) at 37C for 10 minutes. Next, tissue was mechanically disrupted and centrifuged 500g for 5 minutes. Cells were then resuspended in DMEM Ham’s-F12 (LGC^®^) supplemented with 10% FBS (Gibco^®^), 1% penicillin/streptomycin, 1% non-essential amino acids (LGC^®^), 1% sodium pyruvate (LGC^®^) and 1% L-glutamine (LGC^®^) and seeded in cultures flasks maintained overnight at 37°C, 5% CO_2_ atmosphere. Cells were then washed with phosphate saline buffer (PBS) and replaced with fresh media. The media were changed every three days until the confluency was reached, when culture flasks were shaken at 200rpm for 3 hours at 37 C for detaching of microglia. Floating cells were removed, and the remaining attached cells were harvested for *in vitro* assays.

### *In vitro* infection with SARS-CoV2

Primary astrocytes were infected with SARS-CoV-2 at MOI 0.1 for 1h at 37 C and 5% CO_2_ atmosphere. After infection, inoculum was washed, and cells were incubated with fresh DMEM HAMs F-12 supplemented with 10% FBS (LGC^®^), 1% penicillin/streptomycin (LGC^®^), 1% NEAA (LGC^®^), sodium pyruvate (LGC^®^) and L-glutamine (LGC^®^) at 37°C and 5% CO_2_ atmosphere. Supernatant and cells were harvested after 24-, 48- or 72-hours post infection for further analysis. For the experiments with inhibitors astrocytes were treated with inhibitors for 2h prior to infection. For experiments with glucose (17 mM) and glutamine (2mM) supplementation, supplementation was also added 2h prior infection.

### *In vivo* Infection of Golden Syrian Hamsters with SARS-CoV-2

SARS-CoV-2 strain human/BRA/SP02cc/2020 (GenBank access number: MT350282.1) was used in the *in vivo* experiments of this study. This virus was obtained from nasopharyngeal swabs from the first patient (HIAE01) to be diagnosed with COVID-19 in Brazil, isolated in Vero ATCC CCL-81 cells and quantified by using the Median Tissue Culture Infectious Dose (TCID50) assay. This sample was also confirmed to be free of other common 18 human respiratory viruses using a qPCR respiratory panel (Araujo et al. 2020). A third passage aliquot was used to infected hamsters. Brain samples from SARS-CoV-2-infected and uninfected hamsters from two unrelated experiments were used herein (one performed with 15–16-week-old females and another with 18-week-old males). Accordingly, conventional hamsters were acquired from the Gonçalo Muniz Institute, Fiocruz, Salvador, Brazil and maintained at the biosafety level 3 animal facility of the Department of Parasitology, Institute of Biomedical Sciences (ICB), USP, Brazil, with food and water ad libitum. Animals were housed in pairs and groups homogenized based on weight. Animal procedures were approved by the Institutional Animal Care and Use Committees of the ICB and the College of Veterinary Medicine, USP (protocols #9498230321, #3147240820, #2076201020). In both experiments, animals were anesthetized intraperitoneally with 100 mg/kg of ketamine and 5 mg/kg of xylazine at day zero for the intranasal inoculation of 50 l of 1 x 10^5^ TCID_50_ of SARS-CoV-2 suspended in DMEM with 2% FBS (infected group) or 50 μl of sterile DMEM with 2% FBS (uninfected, mock group).

Hamsters were followed daily for clinical signs and weight. Subgroups (n = 3 to 5) of animals were euthanized with a combination of 5 mg/kg of morphine followed by an overdose of ketamine (600mg/kg) and xylazine (30 mg/kg) on days 3, 4, 5, 7 and/or 14 post-infection, depending on the experiment. Brains were collected aseptically at necropsy, using sterile surgical forceps and scissors, and directly placed in 50 mL falcon tubes with DMEM. Brain samples were collected from three infected female hamsters on day 3, and five infected and four uninfected hamsters on day 7. Brain samples were collected from 5 infected male animals on days 3, 5, 7, and 14, and from three non-infected hamsters on day 7.

### Drugs and Inhibitors

This study was conducted using the following drugs and inhibitors: 6-diazo-5-oxo-L-nor- Leucine, 50uM (DON, Cat. Number. 157-03-9; Cayman Chemicals, Ann Arbor, Michigan, USA); 2-deoxy-D-Glucose-6-phosphate, 5mM (2-DG, Cat. Number.17149; Cayman Chemicals, Ann Arbor, Michigan, USA) and Etomoxir, 3uM (Cat. Number. 11969; Cayman Chemicals, Ann Arbor, Michigan, USA).

### Plaque-forming unit assay

For virus titration, Vero CCL81 cells were seeded in 24 well plates one day before the infection for adhesion. Supernatants from *in vitro* assays were serially diluted in pure DMEM low glucose and inoculated for 1 hour at 37°C. Next, the inoculum was removed and replaced with DMEM low glucose containing 2% FBS, 1% penicillin/streptomycin and 1% carboxymethylcellulose. After 3 days, the media was removed, and the cells fixed with 3.7% formaldehyde overnight. The cell monolayers were stained with 1% crystal violet for 15 minutes. The viral titer was calculated based on the count of plaques formed in the wells corresponding to each dilution expressed as plaque-forming units per mL (PFU/mL).

### RNA extraction, viral load, and gene expression analyses

RNA extraction from cell lysates was performed using the MagMAX™ Viral/Pathogen II (MVP II) Nucleic Acid Isolation Kit (Catalog number: A48383) (Applied Biosystems^®^) and carried out according to the manufacturer’s instructions. RNA extraction was performed using Trizol^®^ reagent (ThermoFisher^®^) and carried out according to the manufacturer’s instructions. RNA quantity was determined by NanoDrop (ThermoFisher^®^). Total RNA samples (up to 2 µg) were reverse transcribed using the oligo(dT) primer from the High-Capacity cDNA Reverse Transcription Kit (Thermo Fisher^®^). Molecular detection of SARS-CoV-2 was performed with TaqMan™ Gene Expression Assay (Cat. N. 4331182, ThermoFisher^®^) with specific primers/probes previously described^40^. The quantitative assay was performed using a standard curve produced with serial dilutions of SARS-CoV-2 RNA expressed on a Briggsian logarithm scale as genome equivalents per ug of total RNA. Gene expression assays were performed using Power SYBR® Green Master Mix (Cat. N. 4368577, ThermoFisher^®^) with primers described below. The median cycle threshold (C_t_) value from experimental replicates and 2^-DDCt^ method were used for relative quantification analysis and all C_t_ values were normalized to *Actb.* All qRT-PCR assays were performed on QuantStudio™ 3 Real-Time PCR System (Thermo Fisher^®^).

### Proteomics

#### LC-MS/MS Analysis

After infection hamster astrocytes were resuspended in lysis buffer (100mM Tris HCL, 150 mM NaCl, 1 mM EDTA, 1% Triton X) with freshly added protease and phosphatase inhibitors (Protease Inhibitor Cocktail, SIGMA®) and subjected to ultrasonication (3 cycles of 20s). Samples were submitted to the FASP protocol (Distler U., and Tenzer S., 2016). Samples had their buffer exchanged in a microcolumn tip with a 10kDa MW cut off. Tryptic digestion was performed in column. Forty micrograms of protein were used to carry out the FASP protocol, where the samples were reduced, alkylated, and later digested using trypsin. Digested peptides from each sample were resuspended in 0.1% formaldehyde. The separation of tryptic peptides was performed on an ACQUITY MClass System (Waters Corporation^®^). One μg of each digested sample was loaded onto a Symmetry C18 5 μm × 20 mm precolumn (Waters Corp^®^) used as trapping column and subsequently separated by a 120 min reversed phase gradient at 300 nL/min (linear gradient, 3–55% ACN over 90 min) using a HSS T3 C18 1.8 μm, 75 μm × 150 mm nanoscale and LC column (Waters Corp^®^) maintained at 30 °C. For the gradient elution water-Formic Acid (99.9/0.1, v/v) has been used as eluent A and Acetonitrile Formic Acid (99.9/0.1, v/v) as eluent B. The separated peptides were analyzed by High Definition Synapt G2-Si Mass spectrometer directly coupled to the chromatographic system. Differential protein expression was evaluated with a data-independent acquisition (DIA) of shotgun proteomics analysis by Expression configuration mode (MSe). The mass spectrometer operated in “Expression Mode” switching between low (4 eV) and high (25–60 eV) collision energies on the gas cell, using a scan time of 1.0s per function over 50–2000 m/z. All spectra were acquired in Ion Mobility Mode by applying a wave velocity for the ion separation of 1.000m/s and a transfer wave velocity of 175m/s. The processing of low and elevated energy, added to the data of the reference lock mass ([Glu1]-Fibrinopeptide B Standard, Waters Corp^®^) provides a time-aligned inventory of accurate mass retention time components for both the low and elevated-energy (EMRT, exact mass retention time). Each sample was run in three technical replicates. Continuum LC-MS data from three replicate experiments for each sample have been processed for qualitative and quantitative analysis using the software Progenesis (Waters Corp^®^). The qualitative identification of proteins was obtained by searching in an unreviewed database *Mesocricetus auratus* (Syrian Golden hamster) UNIPROT Proteome ID UP000189706. Search parameters were set as: automatic tolerance for precursor ions and for product ions, minimum 1 fragment ions matched per peptide, minimum 3 fragment ions matched per protein, minimum 1 unique peptide matched per protein, 2 missed cleavage, carbamydomethylation of cysteines as fixed modification and oxidation of methionine as variable modifications, false discovery rate (FDR) of the identification algorithm < 1%. Label free quantitative analysis was obtained using the relative abundance intensity integrated in Progenesis software, using all peptides identified for normalization. The expression analysis was performed considering technical replicates available for each experimental condition following the hypothesis that each group is an independent variable. The protein identifications were based on the detection of more than two fragment ions per peptide, and more than two peptides measured per protein. The list of normalized proteins was screened according to the following criteria: protein identified in at least 70% of the runs from the same sample and only modulated proteins with a p< 0.05 were considered significant.

### Metabolomics of Hamsters astrocytes

Hamster’s primary astrocytes were cultured as described above. Next, the medium was washed twice with phosphate-buffered saline (PBS) at physiologic pH and cells were collected with 600 μL of methanol (MeOH). Finally, the samples were dried and stored in the freezer at -80 °C until metabolites extraction.

#### Bligh-Dyer polar metabolites extraction

Volumes of 150 μL of water (H2O), 190 μL of methanol (MeOH), and 370 μL of chloroform (HCCl 3) were added, and then the tubes were shaken vigorously for 2 minutes. Subsequently, the dimensions were centrifuged for 5 minutes at 13,000 g. The aqueous supernatant was collected and dried for 60 in a concentrator. All samples were stored in a freezer at -80 °C until analyzed by UPLC-MS/MS.

#### Metabolites Analysis

The samples were resuspended in 80 μL of a 1:1 mixture of acetonitrile:water (ACN:H_2_O). For each analysis, we injected 3 μL of sample, and the separation was performed by hydrophobic interaction liquid chromatography (HILIC) analysis using an acquity UPLC^®^ BEH amide 1.7 mm, 2.1 mm x 100 mm column. The mobile phases used for the separations were ACN:H_2_O (80:20) as mobile phase A and ACN: H_2_O (30:70) as mobile phase B, containing 10 mM of ammonium acetate (NH_4_ CHOO) and 0.1% of ammonium hydroxide (NH_4_OH) was additive in both mobile phases. Then, the separation was performed by isocratic flow of 0.3 mL/min, to start with 99% A and was up to 1% A in 7 min. In the final part, we return to 99% A in 1 min and remain for 2 min to equilibrate the column before the next injection. The total run time was 10 min. Negative ion mode was recorded and the instrument was operated in MS E mode in the m/z range of 50–800 Da, with an acquisition time of 0.1 s per scan. The raw files were preprocessed by Progenesis QI Waters® software. The identification was performed with a 5-ppm error for the precursor ion.

### Real-time metabolic assays

An XFe24 Extracellular Flux analyzer (Agilent^®^) was used to determine the bioenergetic profile. Cells were plated at 2 x 10^4^ cells per well in XFe24 plates 24h before infection with SARS-CoV-2. Glycolytic Stress and Mito Stress Tests were performed on XFe24 Bioanalyzer at 72 hpi. All assays were performed following manufacturer’s protocols. Results were normalized to cell number.

### Electron microscopy

Hamster astrocytes infected and uninfected were recovered, washed with PBS 1x and fixed with Glutaraldehyde 4% overnight. Samples were washed with PBS 1x and followed to dehydration process with acetone gradient, desiccation, and gold metallization. Finally, samples were analyzed in JOEL1010 transmission electron microscope. Generated images were loaded into Fiji software for further analysis. For TEM mitochondrial measurements, the images were filtered and manually measured for width and length.

### Mitochondrial analysis

For measurements of mitochondrial superoxide, the cells were stained with LIVE/DEAD™ Fixable Green (L34970; ThermoFisher^®^, 15 min) and MitoSOX™ Red mitochondrial superoxide indicator (M36008; ThermoFisher^®^; 2.5uM, 10 min). Cells were then washed with warm 1X PBS and fixed with 4% PFA for 15 minutes at 4C. Next, cells were analyzed using a Accuri C6 Plus (BD Biosciences^®^) cytometer and data analyzed using FlowJo X software.

### Immunofluorescence

On day 7 of astrocyte culture, 2 x 10^4^ cell were seeded in a 24-well plate, after overnight incubation with culture media cells were fixed with ice-cold 4% paraformaldehyde in 0.1 M phosphate buffer pH 7.4 for 30 minutes. After antigen retrieval (Tris/EDTA-Tween-20 buffer -10mM Tris, 1mM EDTA, 0.05% Tween 20, pH 9 - for 30 minutes at 95°C, followed by 20 minutes at room temperature), cells were permeabilized with 0.5% Triton-X100 in PBS for 10 minutes, blocked with blocking solution (5% donkey serum + 0.05% Triton X-100 in PBS) (Sigma-Aldrich) for 2 hours at room temperature and incubated with primary antibodies: GFAP (1:1000, Sigma-Aldrich), IBA1 (1:500, Abcam), and MAP2 (1:500, Sigma-Aldrich) diluted in blocking solution, overnight at 4°C. The cells were PBS washed and incubated with secondary antibodies (AlexaFluor 594; AlexaFluor 488; 1:2000, Invitrogen) diluted in PBS + 0.05% Triton X-100 for 2 hours at room temperature, protected from light. For nuclear staining, cells were incubated for 20 minutes at room temperature with DAPI (4&#39;,6-diamidino-2-phenylindole; 1:150000, Sigma-Aldrich), mounted on glass slides with Fluoromount-G (Invitrogen) mounting medium and sealed with nail polish. Images were acquired using Nikon Eclipse 80i fluorescence microscope (Nikon Instruments Inc., NY, USA) coupled with Nikon Digital Camera DXM 1200 C. Quantification was made using ImageJ software (NIH), where 5 to 10 randomly chosen visual fields per coverslip from three independent cultures were analyzed.

### Hamster brain slice culture and SARS-CoV-2 infection

Brain sections were obtained from male hamsters and prepared as previously described, with modifications (Mendes et al., 2018; Fernandes et al., 2019). Whole brain - cerebellum and brainstem were removed - was sliced at 200 µm in a VT1000s automatic vibratome (Leica) and cultivated free-floating with Neurobasal A (Gibco) medium supplemented with 1% Glutamax (Gibco), 1% penicillin/streptomycin (Gibco), 2% B27 (Gibco), and 0.25µg/mL fungizone (Sigma). For virus infection, the medium was removed, and the brain slices were exposed to 3x10^6^ TCID50 of SARS-CoV-2 or an equivalent volume of mock medium. Infection was performed for 2h at 37°C and 5% CO2 in a biosafety level 3 laboratory. The inoculum was removed, the tissue was washed, and the slices were maintained in fresh medium at 37°C and 5% CO2 until processing for subsequent analysis. This procedure was approved by the CEUA- FMRP (#066/2020).

### Single Nuclei Transcriptomic Profile Analysis

Raw snRNA-seq data were obtained from frozen medial frontal cortex tissue from six post-mortem control and seven COVID-19 patients assessed herein were obtained from the dataset GSE159812 on NCBI GeneExpression Omnibus public databank. Reads were aligned to the hg38 genome (refdata-gex-GRCh38-2020-A) using CellRanger software (v.6.0.0) (10x Genomics) generating the raw gene counts matrix. Reads mapping to pre-mRNA were also counted to account for unspliced nuclear transcripts. After quantification using the CellRanger count pipeline on each individual library, the CellRanger aggr pipeline was used to aggregate the libraries into two distinct groups, control and infected. We subsequently used these grouped matrices as input to downstream analysis in Seurat (version 4.0.4)^29^, initially filtering nuclei with unique molecular identifiers over 2,500 or less than 200, besides outliers with more than 5% mitochondrial counts. We performed normalization regressing out heterogeneity associated with mitochondrial contamination fitting the counts to a Gamma-Poisson General Linear Model, removing this confounding source of variation through scTransform Seurat’s function^30^ and glmGamPoi package^31^. Integration of both groups was executed identifying cross-dataset pairs of cells that are in a matched biological state (‘anchors’), correcting technical differences between those groups to improve comparative snRNA-seq analysis. Dimensionality reduction was implemented by Principal Component Analysis, followed by UMAP embedding using the first 20 principal components, the same parameter as used for clustering with both FindNeighbors and FindClusters (at 0.5 resolution) Seurat’s functions. Marker genes were identified through MAST positive differential expression^32^ of each cluster against all others and cell types annotation was achieved using previously published marker genes^33^ . Differential gene-expression comparison between the control individuals and infected patients was done using the MAST algorithm (v.1.18.0). Genes with fold change > 0.25 (absolute value), adjusted P value (Bonferroni correction) < 0.1 and detected in a minimum fraction of 10% nuclei in either of the two populations were considered differentially expressed. Differentially Expressed Genes (DEGs) corresponding to the Differentially Expressed Proteins (DEPs) from hamster brain proteomics and concordant fold-change were subjected to Gene Ontology analysis, performed through online database PANTHER using PANTHER Pathways dataset^34^.

### Statistical analysis

Data was plotted and analyzed using the GraphPad Prism 8.0 software (GraphPad Software^®^, San Diego, CA). For analyses between 2 groups, Student’s *t* test was used. For comparisons among 3 or more groups, *One*-way ANOVA, followed by Tukey post hoc tests was used as described in figures legends. Differences were considered statistically significant when P value was <0.05.

## RESULTS

### SARS-CoV-2 infects Syrian hamsters’ astrocytes

We started our experiments by infecting brain slices of adult Syrian golden hasmters with 3x10^6^ TCID_50_ of SARS-CoV-2 (Supplementary figure1A). Tissue infection was confirmed by spike protein detection in GFAP^+^ astrocytes (Supplementary figure 1B) and corroborated by increased nucleocapsid RNA (N1) concentration was detected 48 hpi (Supplementary Figure 1C). Thus, we decided to specifically investigate the impact of SARS-CoV-2 infection on astrocytes. For that, hamster’s primary cultures obtained from 2-4 days old pups composed of more than 80% astrocytes and very few neurons and microglia (Figure 1A left panel – bar graph) were infected with MOI 0.1 of SARS-CoV-2 (strain SPPU-0691/B.1.1.28 Genbank: MW441768#). Confirming data from tissue slices, increased amounts of SARS-CoV-2 RNA were detected in the cell cultures at 48- and 72-hours post infection (hpi) (Figure 1B), confirmed by viral particles detected in the culture supernatant in which the viral load was significantly increased at 72 hpi compared to 24 hpi for both MOIs tested (Figure 1C). To further confirm the presence of viral particles in cell cultures, we performed transmission electron microscopy (TEM) for viral particles visualization. Figure 1D shows viral particles attached to the cell surface (black arrowheads), as well as the presence of double-membrane vesicles (asterisks), characteristics of coronavirus infection^35^.

**Figure 1.**
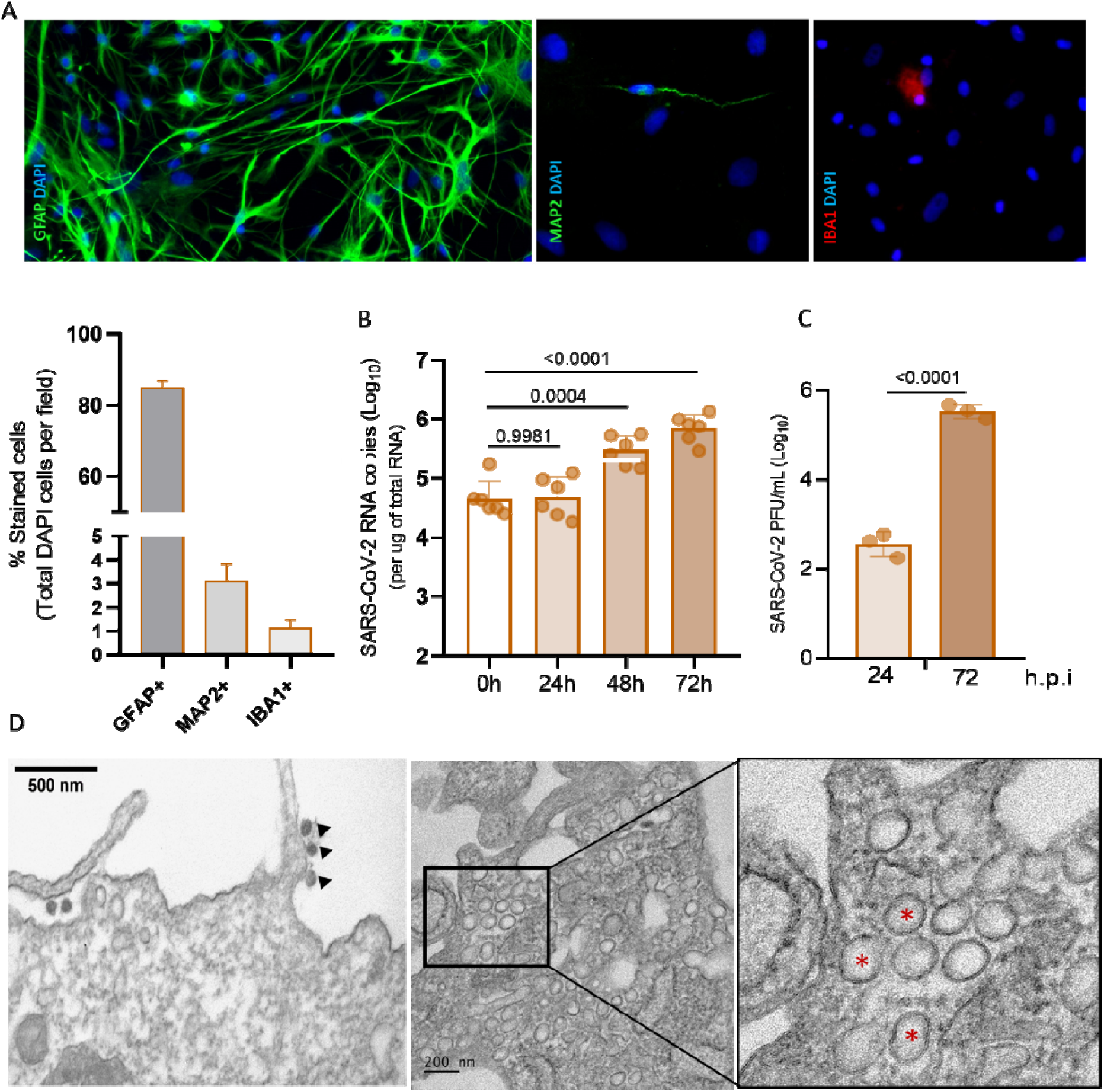
SARS-CoV-2 infects hamsters’ astrocytes in vitro. Immunofluorescence of primary astrocytes stained for GFAP (green), IBA-1 (red) and MAP-2 (green) for culture characterization. Results expressed as percentage of immunoreactive cells for each cell type (GFAP-astrocytes, IBA1 – microglia, MAP2 – neurons) over the total number of cells labeled with DAPI per field ± SEM. Minimum of 5 to 10 fields per coverslip from three independent experiments were analyzed. Scale bar = 50µm (A). Viral RNA (B) and infectious particles (C) were detected in cultures infected with SARS-CoV-2 at MOI 0.1 by qPCR and PFU assay, respectively. Statistical analyses were performed by multiple *t*-tests and p < 0.05 was considered significant. Transmission electron microscopy of astrocytes infected with MOI 0.1 at 72hpi (D). Black arrows indicate SARS-CoV-2 particles attached to the cell surface and red asterisks indicates double-membrane vesicles. Cultures performed in triplicates and graphs representative of two experiments.

**Table 1.**
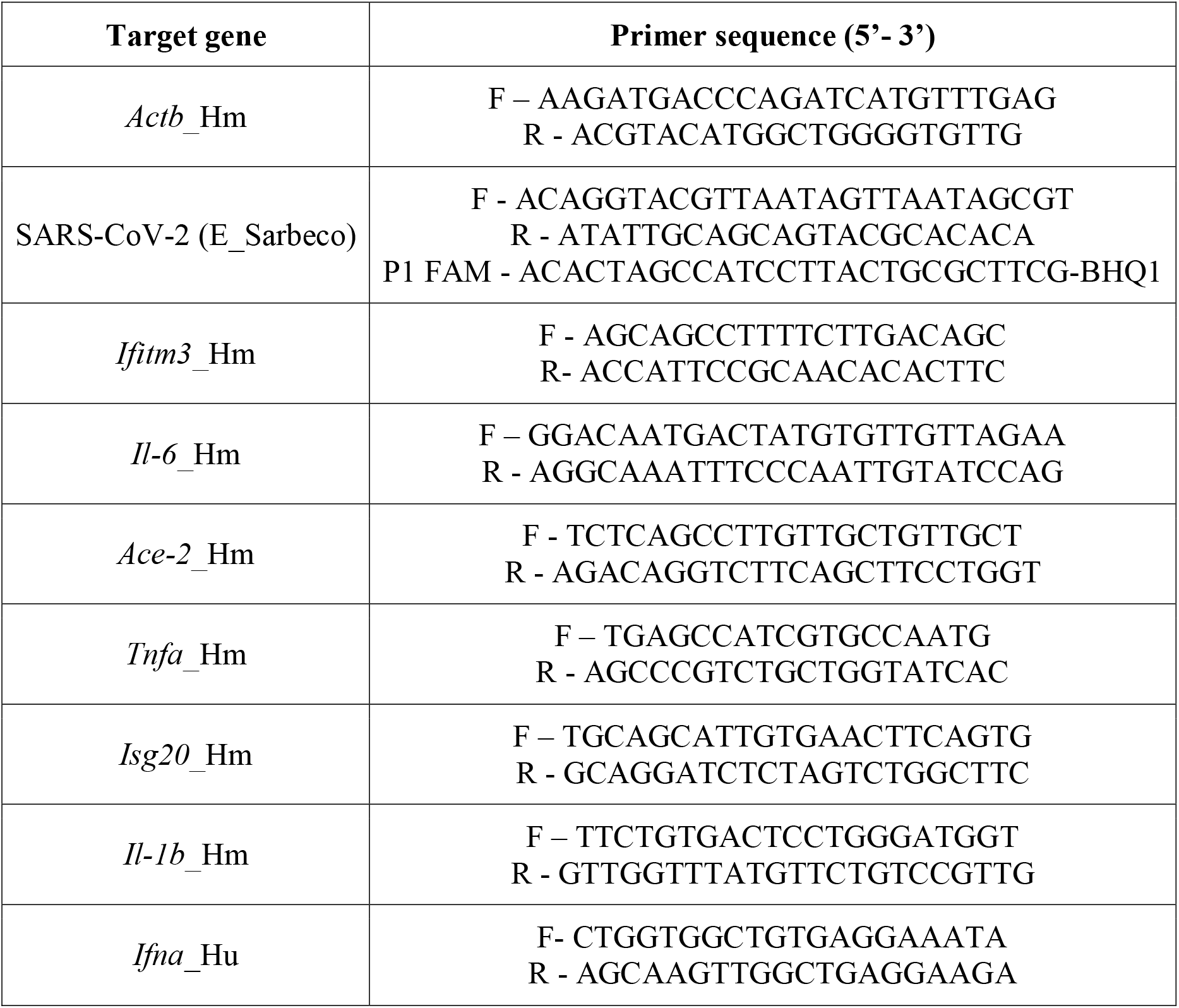
Primers used for gene expression analysis.

### SARS-CoV-2 astrocyte infection induces pro-inflammatory cytokines, interferon-stimulated genes and changes the expression of carbon metabolism related proteins *in vitro*

Next, we evaluated whether SARS-CoV-2 triggers the production of pro-inflammatory cytokines in hamster primary astrocytes. Consistent with the literature, we observed that *Il-1b*, *Il- 6*, *Tnf-a* and *Ifn-a* are significantly increased 72 hpi at MOI 0.1 (Figure 2A-D). Confirming that type I interferons are active, we also observed that the interferon-stimulated genes (ISGs)*, Isg20, Iftm3* and including *Ace-2,* were up-regulated (Figure 2 E-G).

**Figure 2.**
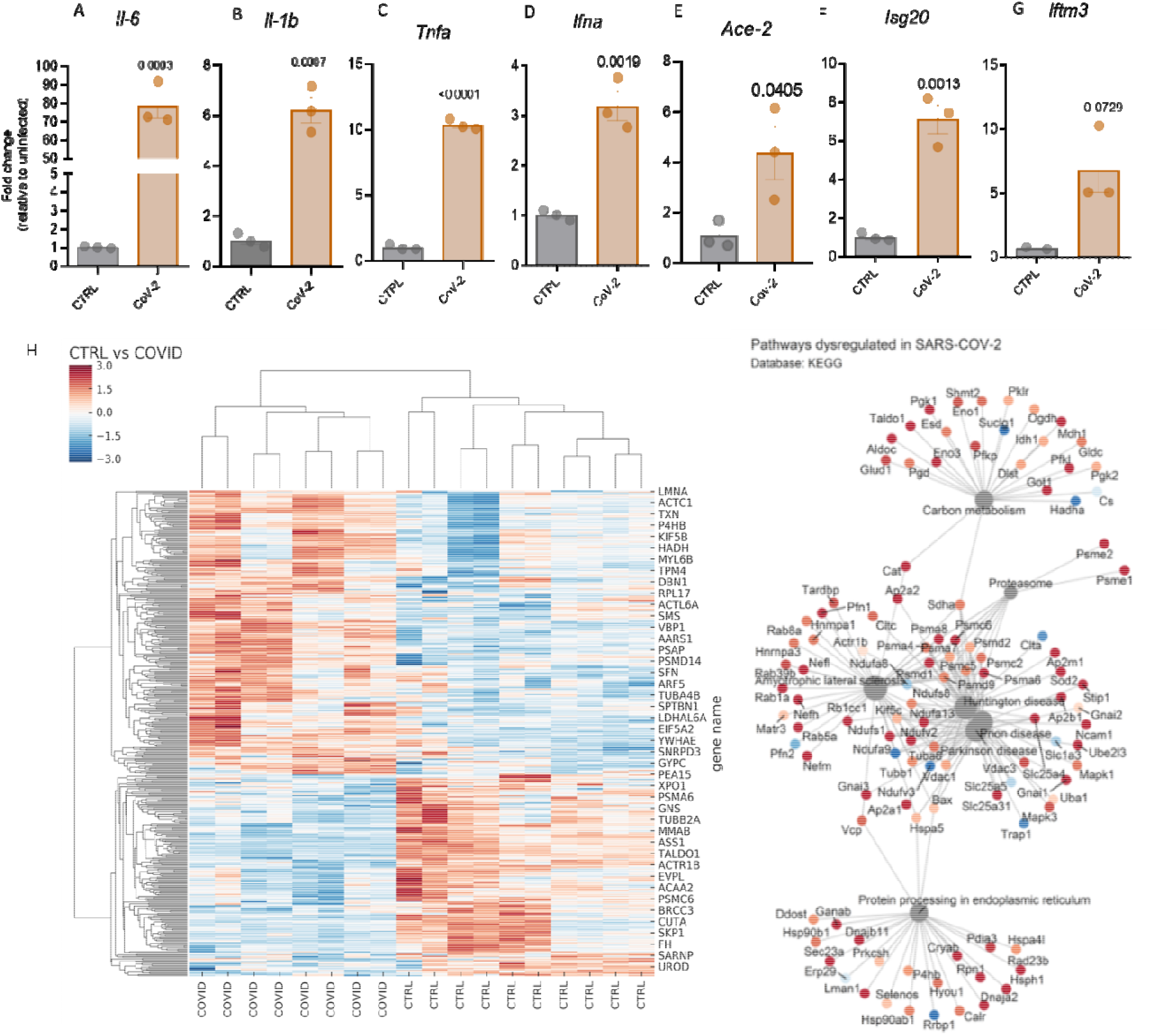
SARS-CoV-2 induces pro-inflammatory cytokines and interferon stimulated genes expression hamsters’ primary astrocytes. Quantitative PCR for pro-inflammatory (**A-D**) and interferon stimulated genes (**E-G**) of cultures infected with SARS-CoV-2 MOI 0.1. The fold change normalized by the relative expression of control cultures. Statistical analyses were performed by *t*-test and p < 0.05 was considered significant. Heatmap of proteomic analysis from cultures infected with SARS-CoV-2 MOI 0.1. n=8 for infected and n=10 for control samples (H). Colors represents the protein z-score. DEPs from proteomic data submitted to KEGG database analysis (**I**). The nodes in gray represent major pathways and the color of the genes the z-score (red to blue). Cultures performed in triplicates and graphs representative of two experiments.

Next, we sought to evaluate the overall impact of SARS-CoV-2 infection on the protein expression of infected astrocytes. We infected the cell cultures with MOI 0.1 and performed proteomic analysis at 72 hpi. We observed the differential expression of 646 proteins (DEP), of which 568 were up-regulated and 78 down-regulated in infected cells compared to controls (Figure 2H and Supplementary Figure 2A). A pathway-protein analysis using KEGG database evidenced that the great majority of DEP correlate to carbon metabolism (Got1, Eno3, Aldoc, Tald1, PgK1, Ogdh, Pkfp, Pfk1, Gfpt), tricarboxylic acid cycle (Mdh1, Acly, Sdha, Ogdh, Dlst, Idh1, Cs, Suclg1), glycolysis (Aldoc, Gapdhs, Pgk1, Ldhb, Pfkl, Eno3, Pfkp, Eno1, Pgk2, Pklr, Aldh9a1), as well as to protein processing in the endoplasmic reticulum (Pdia3, Cryab, Dnajb11, Sec23a, Lman1, Hsph1) (Figure 2K). Of note, these proteins are enriched in pathways that are also dysregulated in several brain diseases, such as Huntington Disease (HD), Parkinson’s Disease and Amyotrophic Lateral Sclerosis (ALS) (Nefh, Rab1a, Rab39b, Pfn1, Psma4, Psmc6, Ndufv2, Gnai3, Ap2a1, Slc25a5) (Figure 2I and Supplementary Figure 2B). Conversely, by comparing our data with data sets from human brain samples from SARS-CoV-2 infected subjects, we observed an overlap of approximately 53 proteins (Supplementary Figure 2C). Consistently, most of these changes are related to alterations in the overall metabolic profile of the cells, including carbon metabolism, glycolysis/gluconeogenesis, and pentose phosphate pathway (Supplementary Figure 2D).

### SARS-CoV-2 induces metabolic changes in astrocytes *in vitro*

Because SARS-CoV-2 infection induced important changes in carbon metabolism-related proteins, we next validated these findings. First, we infected the cultures with SARS-CoV-2 (MOI 0.1) and assessed mitochondrial morphology through transmission electron microscopy (TEM). As shown in Figure 3A, mitochondria from SARS-CoV-2 samples are more fragmented than in control cultures. This was associated with increased mitochondrial reactive oxygen species (ROS) production at 24 and 72 hpi (Figure 3B). Next, we performed high resolution real-time respirometry at 72 hpi. Conversely, we observed that both glycolytic and non-glycolytic acidification were increased in the presence of the virus (Figure 3C), although, basal and maximal mitochondrial respiration were not statistically different, despite the trend (Figure 3D).

**Figure 3.**
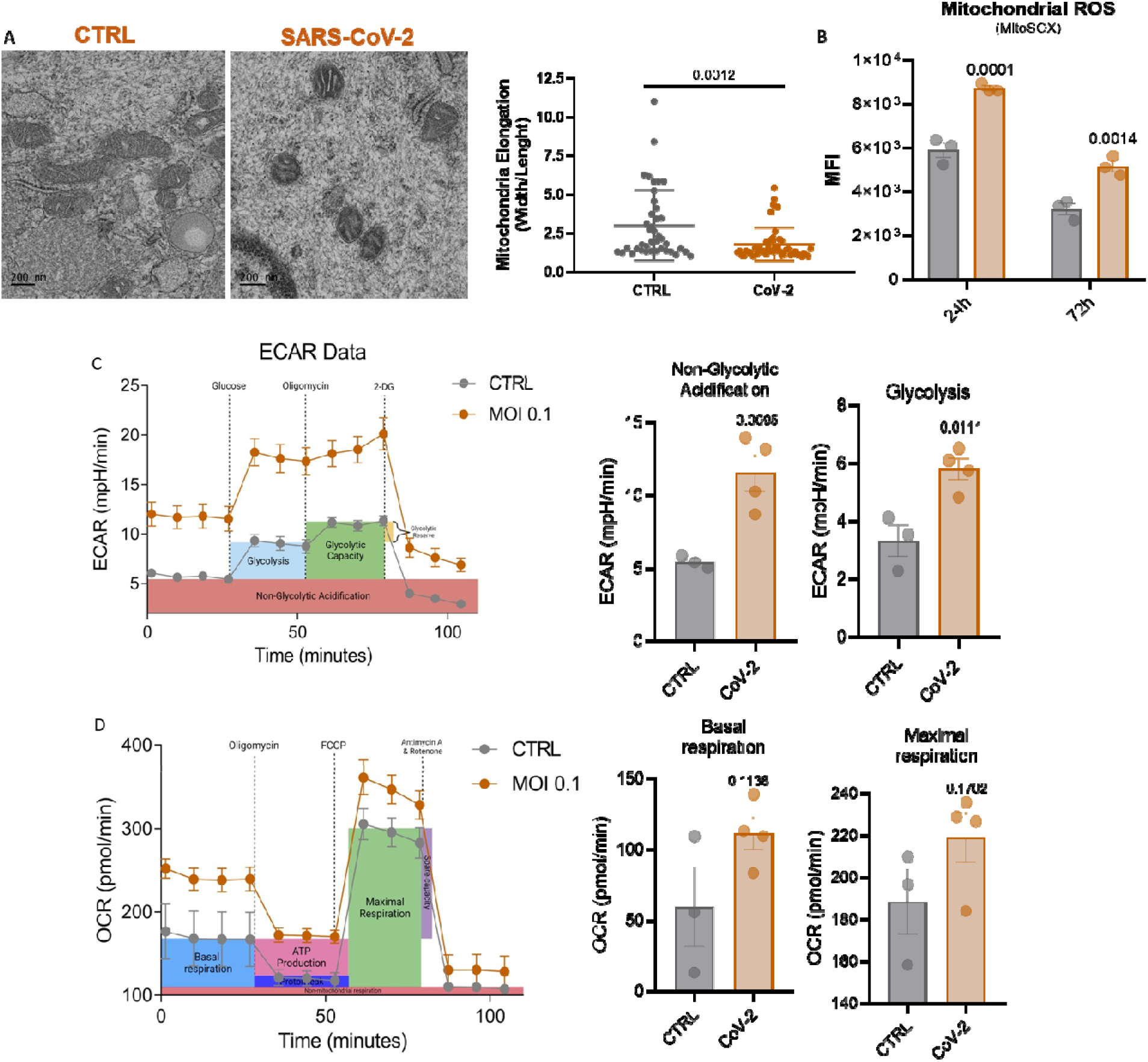
SARS-CoV-2 infection alters mitochondrial morphology, reactive oxygen species production and cellular bioenergetics of hamsters’ astrocytes. Mitochondrial related analysis were performed after SARS-CoV-2 infection (MOI 0.1) at 72 hpi. Mitochondrial morphological changes were evidenced by TEM (scale bar 200nM) (**A**). Statistical analyses were performed by *t*-test. Mitochondrial ROS production was measured by flow cytometry at 24- and 72 hpi, graphs represent median intensity fluorescence (MFI) (**B**). Statistical analyses were performed by *two-* way ANOVA and Sydak’s multiple comparisons test. Bioenergetic profile by oxygen consumption rate (OCR) (**C**) and extracellular acidification rate (ECAR) (**D**). Statistical analyses were performed by *t*-test. Statistical significance: p < 0.05. Cultures performed in triplicates and graphs representative of two experiments.

### SARS-CoV-2 changes the metabolic profile of hamsters’ primary astrocytes

To deeply characterize the metabolic changes induced by SARS-CoV-2 infection in astrocytes, we performed a targeted metabolomic analysis at 72 hpi with MOI 0.1. We observed a significant reduction in almost all metabolites evaluated (Figure 4). The decreased levels of TCA cycle precursors such as lactate (Figure 4A), pyruvate (Figure 4B), serine (Figure 4C), acetate (Figure 4D), aspartate (Figure 4I) 2-hydroxyglutarate (Figure 4L) as well as the intermediates alpha-ketoglutarate (Figure 4J) and succinate (Figure 4K), strongly suggests cataplerotic reactions to supply the synthesis of biomolecules, which may be involving in supporting viral replication. Interestingly, although glutamate levels were maintained, there was a significant decrease of glutamine (Figure 4F), alpha-ketoglutarate (Figure 4J) and 2- hydroxyglutarate (Figure 4L) levels on infected samples. The levels of GABA (Figure 4H), another glutamate precursor, was unchanged. Interestingly, palmitate levels were increased (Figure 4E), further suggesting that TCA intermediates are being used as building blocks for novel biomolecules, such as lipids. Altogether, our data show an intense activation of cataplerotic and anaplerotic pathways, especially with a switch to glutamine and alfa- ketoglutarate consumption and lipid synthesis.

**Figure 4.**
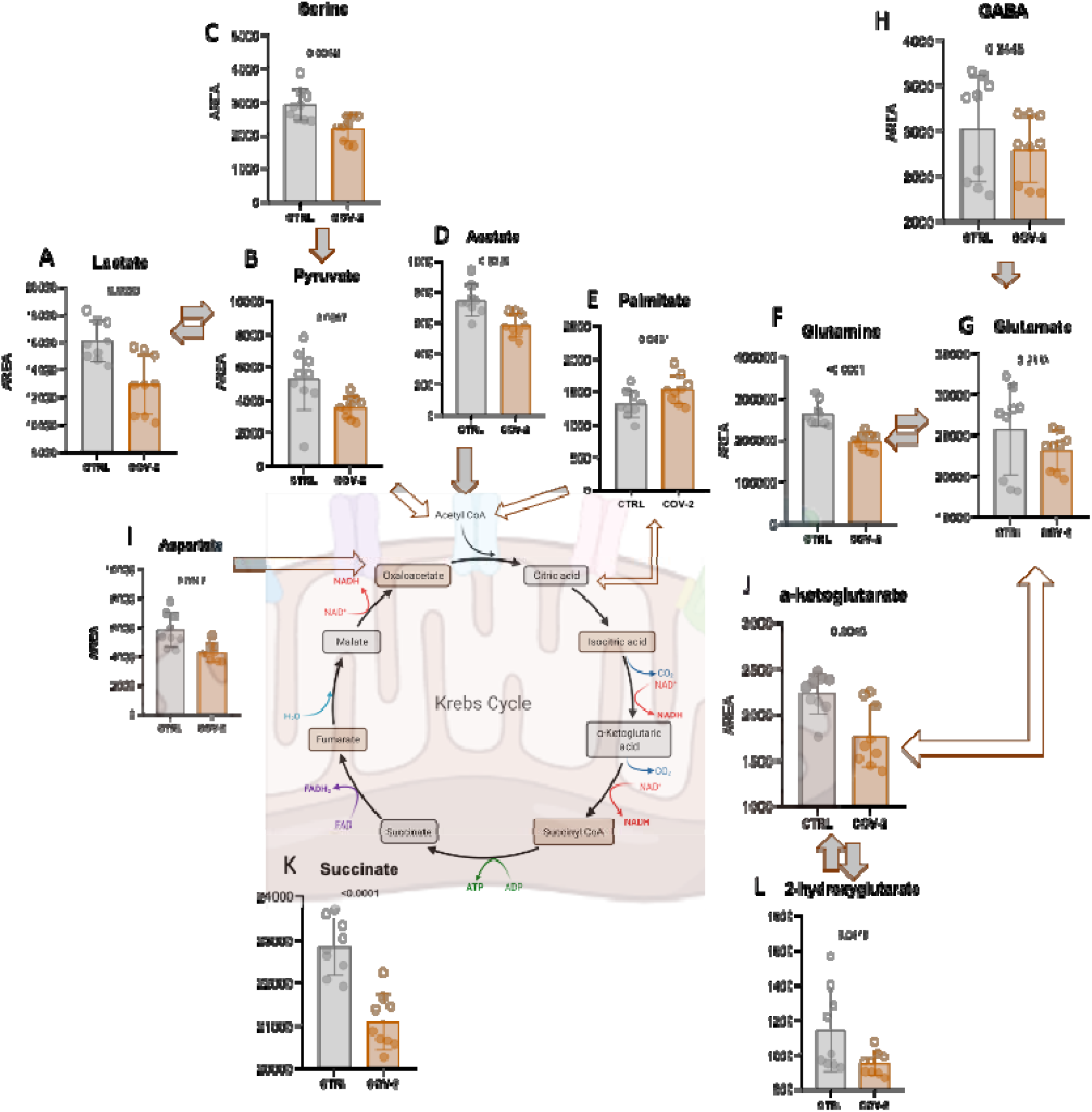
SARS-CoV-2 impacts hamsters’ astrocytes metabolic pathways. Targeted metabolomic data of amino-acids (**C, G, I**), TCA cycle (**A-B, D** and **K**), fatty acid (**E**) and glutamate related metabolites (**F-H, J** and **L**) of hamsters’ astrocytes cultures infected with SARS-CoV-2 MOI 0.1. The arrows represent the relationship between metabolites. The analysis was conducted at 72-hour post infection (MOI 0.1) and data is represented by the area under the curve of each metabolite detection peak. Statistical analyses were performed by Welch’s *t*-test and p < 0.05 was considered significant. Cultures performed in triplicates and graphs representative of two experiments.

### SARS-CoV-2 replication in astrocytes is dependent on glutaminolysis

To further determine which specific metabolic pathway is more important in favoring SARS-CoV-2 replication, we treated infected cultures with inhibitors of fatty acid metabolism (Etomoxir, 3uM), glutaminolysis (L-DON, 50uM) and glycolytic pathways (2-DG, 5mM) as illustrated in Figure 5A. As shown in Figure 5B, blockade of glycolysis with 2-deoxyglucose (2- DG) did not significantly change SARS-CoV-2 replication at 24-72 hpi., further corroborated by PFU assay (Figure 5B right). Also, the use of etomoxir, an inhibitor of the carnitine palmitoyltransferase 1a (CPT-1a), indicates low dependence of mitochondrial fatty acid oxidation (Figure 5B). Interestingly, however, the inhibition of mitochondrial glutaminase (GLS) with L-6-Diazo-5-oxo-norleucine (L-DON), significantly decreased SARS-CoV-2 replication (Figure 5B) and the release of viral particles (Figure 5C). Accordingly, the addition of glutamine (2mM) in the absence of glucose significantly increased viral replication, whereas glucose (17mM) was not able to do so (Figure 5C).

**Figure 5.**
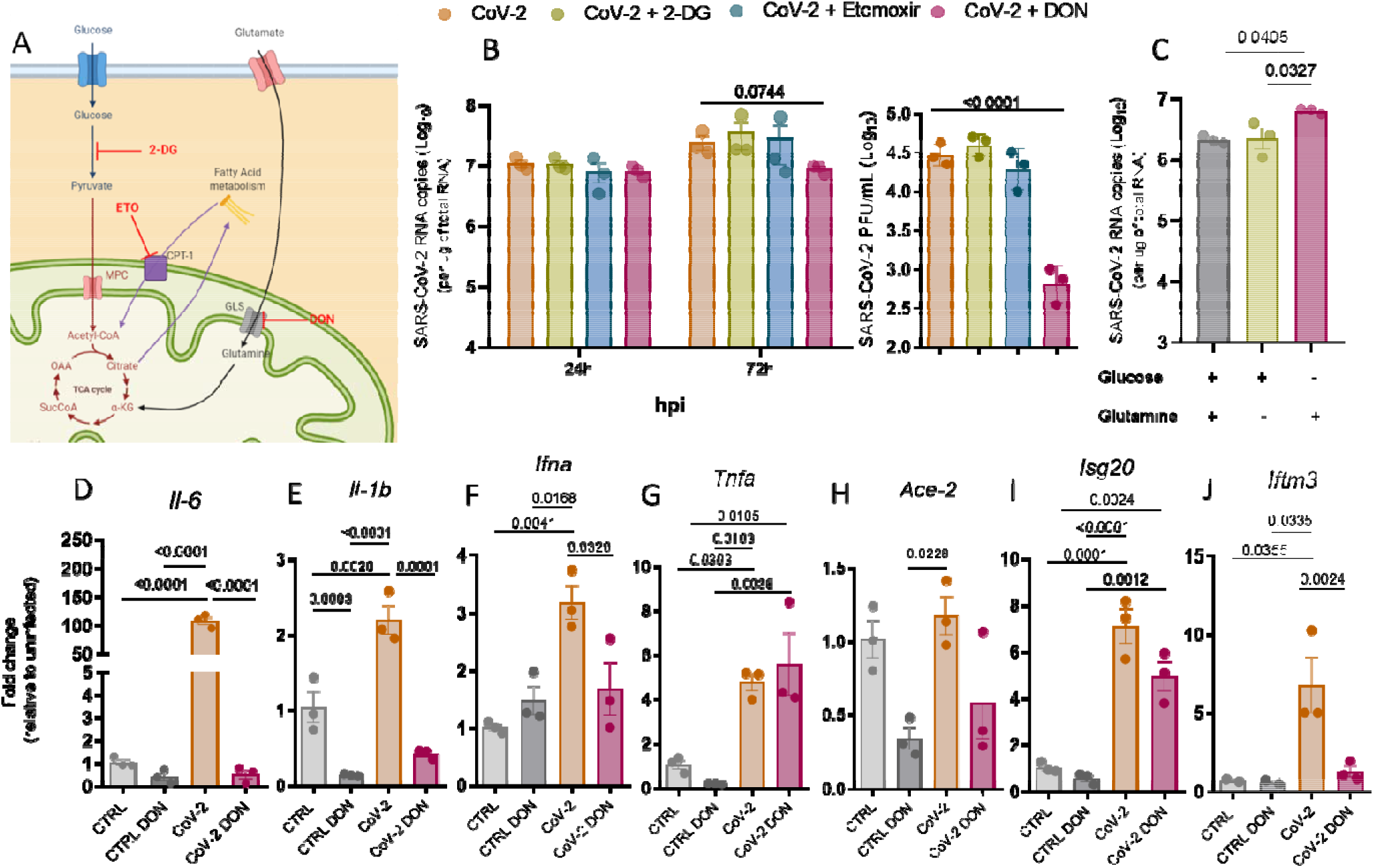
Blockade of glutaminolysis reduces SARS-CoV-2 replication and pro-inflammatory gene expression. Illustration showing the targets of 2-DG (2mM), Etomoxir (3uM) and DON (50uM), blocking glycolysis, β-oxidation and glutaminolysis, respectively (A). Cultures were pre-treated with inhibitors 2 hours prior to SARS-CoV-2 infection (MOI 0.1). Viral RNA and infectious particles were quantified after 72 hpi by qPCR (left) and PFU (right), respectively (**B**). Viral RNA of cultures supplemented or not with glucose (17mM) or glutamine (2 mM) qPCR of pro-inflammatory and interferon stimulated genes (**E-J**). Statistical analyses were performed by multiple t-test (**A**) or one-way ANOVA with Tukey’s multiple comparisons (**B-J**). Cultures performed in triplicates and graphs representative of two experiments.

Next, we evaluated the expression of proinflammatory cytokines during the treatment with L-DON. Blockade of GLS with L-DON decreased the transcription of *Il-1b*, *Il-6* (Figure 5D-E), and *Ifn-a* (Figure 5F) but not *Tnf-a* (Figure 5G) when compared to non-treated groups. This is in accordance with the reduction of the ISGs, *Ace-2, Isg20* and *Iftm3* (Figure 5H-J). These results evidence that glutamine is important for viral replication and the blockade of glutaminolysis reduces viral loads which in turn reduces type I IFN and pro-inflammatory responses

### SARS-CoV-2 RNA is detected in the cortex, hippocampus, and olfactory bulb of hamsters *in vivo*

It has already been demonstrated that the Syrian hamster is a permissive model to nasal infection with SARS-CoV-2^36^. Thus, we decided to use this model to address whether the virus reaches important areas of the brain, such as cortex, hippocampus, and olfactory bulb after intranasal infection with 10^5^ TCID_50,_ as illustrated. Consistently, we confirmed the presence of SARS-CoV-2 in these areas, both at 3-, 5-, 7- and 14-days post-infection (dpi) (Figure 6A) with higher titers at day 3. We also observed increased expression of the same pro-inflammatory cytokines evaluated *in vitro*, *Il-6, Il-1b* and *Tnfa*, as well as the ISGs *Isg20* and *Iftm3*, both in the hippocampus (Figure 6B) and cortex (Figure 6D). Interestingly, although *Il-1b* was increased in the cortex (Figure 6D), no difference was observed in the hippocampus (Figure 6B) the same was observed for *Il-6* which is increased the hippocampus (Figure 6B) but not in the cortex (Figure 6D). As expected, proteomic analysis of these brain samples showed that in the hippocampus, DEPs correlated with pathways of oxidative phosphorylation, long-term potentiation, dopaminergic and cholinergic synapses, among others (Figure 6C). In the cortex, DEPs correlated with important neuronal function, as synaptic vesicle cycle, dopaminergic and GABAergic synapses, as well as to neurological features of brain diseases, as Parkinson’s disease and long-term depression (Figure 6E). To further confirm whether these changes are in accordance with those observed in human COVID-19 patients, we compared the DEPs from our *in vivo* proteomic analysis with a data set from the literature that analyzed more than 64,000 single cells obtained from the frontal cortex and choroid plexus of 8 deceased COVID-19 patients (Figure 6F). We found a strong convergence of important signaling pathways enriched in both datasets, such as those for metabotropic and ionotropic glutamate receptors, Wnt signalling and adrenergic activation. Also, important physiological processes were related between hamster and human samples, as synaptic vessel trafficking, glycolysis, and inflammation, which were further consistent with the enrichment of DEPs correlated to brain diseases, as Parkinson’s disease, Huntington’s disease, and Alzheimer’s disease (Figure 6G). We also performed qPCR for the expression of glutamate transporters and receptors in the hippocampus and cortex of the infected animals, but we detected no differences (Supplementary Figure 4). Altogether, our data support the hypothesis that important changes in protein expression and carbon metabolism is occurring in the brains of infected Golden Syrian hamsters, and these changes may share similarities to those observed in COVID-19 patients.

**Figure 6.**
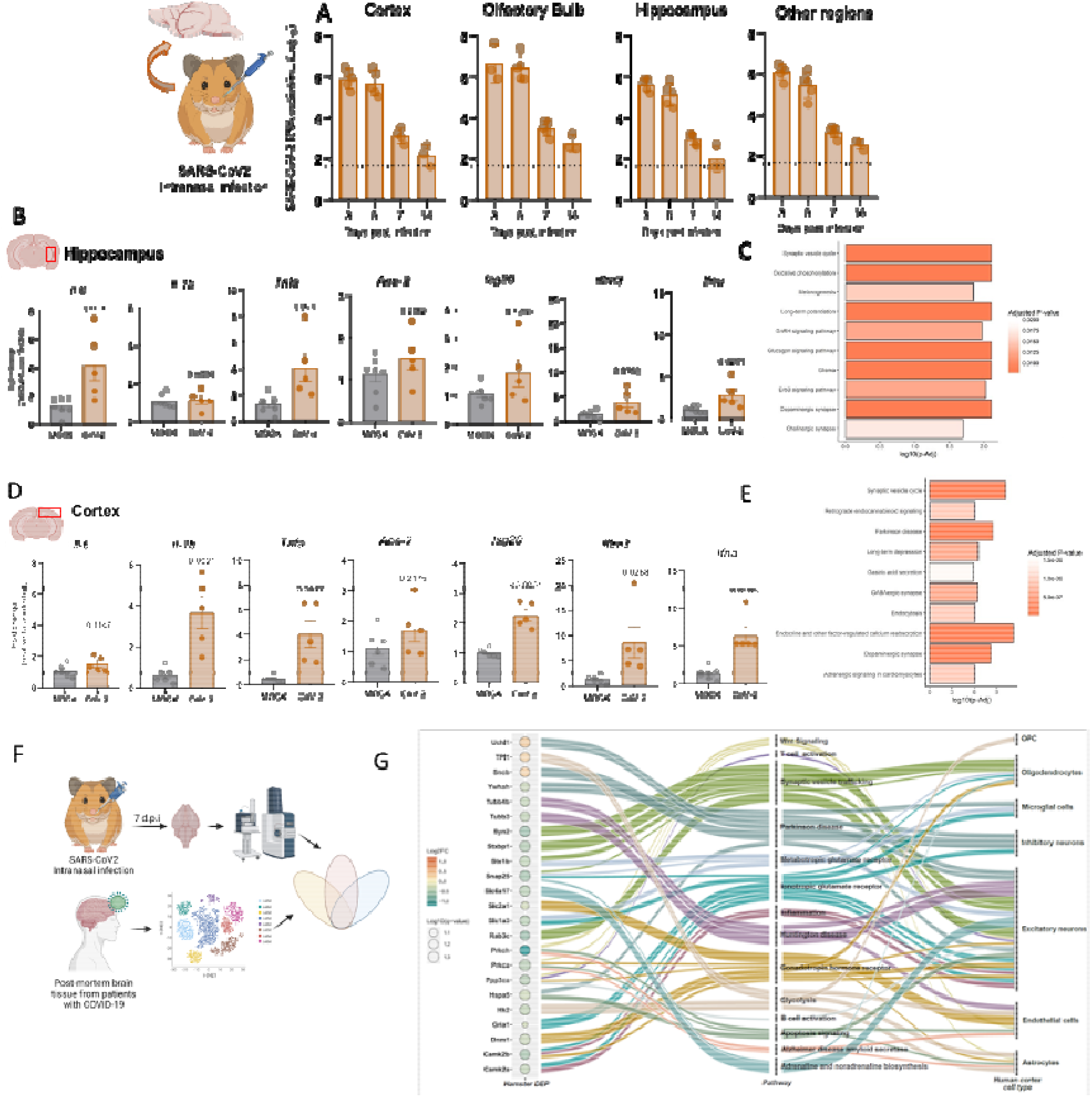
SARS-CoV-2 infects Syrian hamsters’ brains *in vivo* and elicit changes similar to human COVID-19. Animals were inoculated intranasally with 1x10^5^ TCID of SARS-CoV-2 and viral RNA detection was performed by qPCR in different areas of the brain over the infection course (**A**). Cytokine expression by qPCR (**B** and **D**) and KEGG pathways analysis (**C** and **E**) of mock or infected hippocampus and cortex, respectively at 72 dpi. The fold change was normalized by the relative expression of uninfected groups and only significant DEPs (FDR adjusted) were included in the KEGG analysis. Statistical analysis in gene expression graphs were performed by *t*-test and p<0.05 was considered significant. Illustrative scheme of data generation from the comparison of our hamster proteomic data to human data set (**F**). The generated data were fed into an alluvial graph connecting the DEPs from hamster brain proteomics with significant ontology pathways of public human brain single-cell sequencing data sorted by specific brain cell populations (**G**). *OPC = Oligodendrocyte progenitor cell*. Graphs are representative of two experiments.

**Figure 7.**
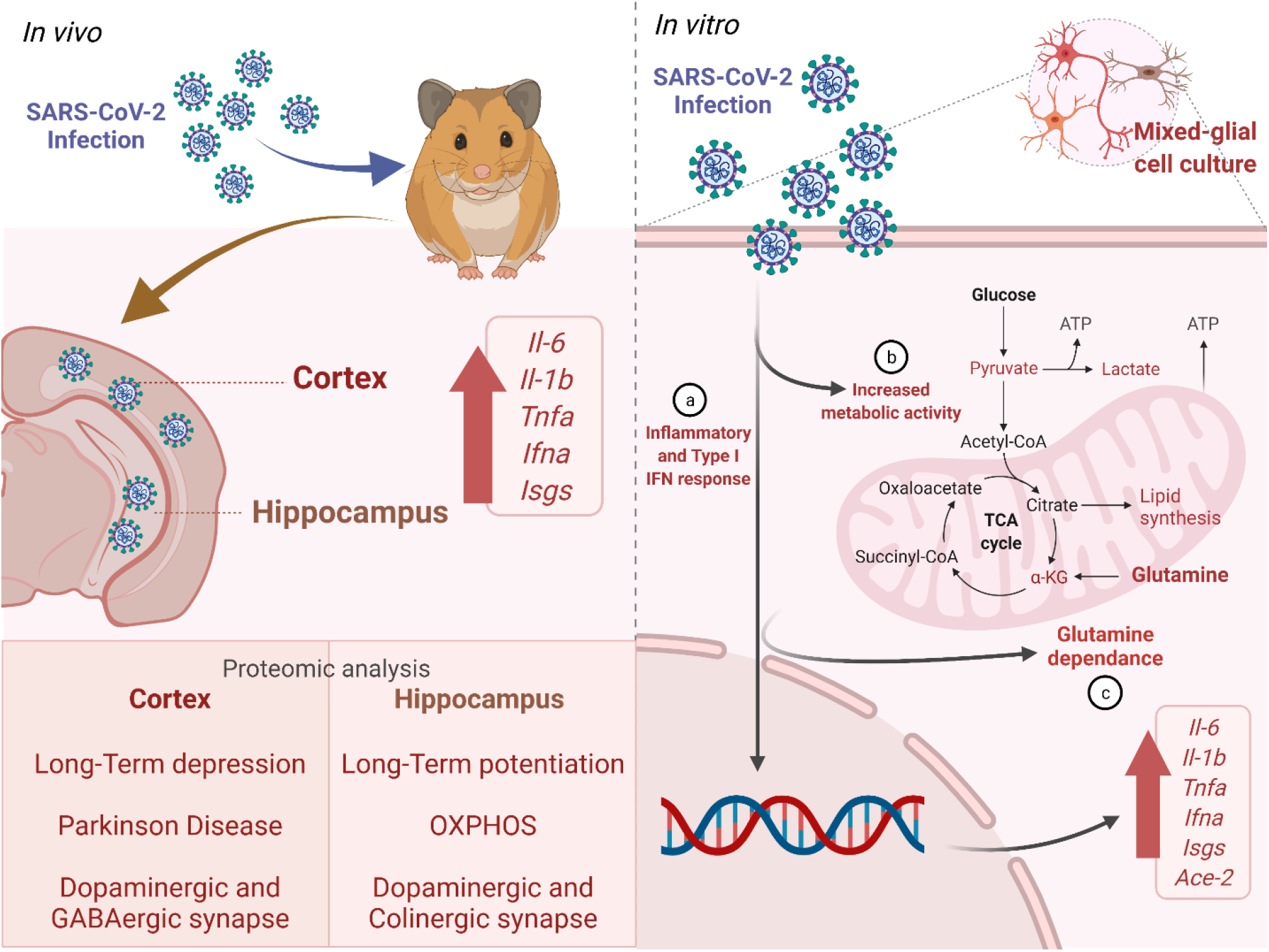
Graphical abstract. SARS-CoV-2 infection of Syrian hamsters’ brain impacts carbon metabolism and protein profile: a glimpse for SARS-CoV-2 induced neurological impact? The presence of SARS-CoV-2 within different regions (olfactory bulb, hippocampus and cortex) of infected Syrian hamster’s brain demonstrates that the virus can avidly reach the central nervous system. The presence of SARS-CoV-2 was associated with increased expression of pro-inflammatory cytokines (*Il-6*, *Il-1b*, *Tnf-a*) and interferon stimulated genes (*Ace-2*, *Isg20*, *Iftm3*) (**a**). Primary mixed glial cells infected *in vitro* also demonstrated dysregulated metabolic pathways such as mitochondrial respiration and glycolysis associated with an important decrease of intermediates and substrates from tricarboxylic cycle (TCA) (**b**). Importantly, the blockage of glutaminolysis was critical for the maintenance of viral replication (**c**). Finally, protein analysis evidenced that SARS-CoV-2 infection results in enriched pathways associated with neurological impacts as long-term depression, memory decline, and Parkinson’s disease, which are also found in single cell analysis of COVID-19 patients’ brains. This figure was designed with Biorender.com

## DISCUSSION

Although the great majority of people infected with SARS-CoV-2 develop only mild or asymptomatic disease, COVID-19 has already taken more than 5.7 million lives worldwide. Moreover, although it is essentially a lung disease, unconventional neurological symptoms have been reported since the beginning of the pandemic, from mild anosmia and ageusia^19, 21^, to more severe neurological cases of encephalitis, cerebrovascular disease, ataxia and seizures^37^. Most recently, the description of the “long COVID-19” and the “brain fog”, still intrigues physicians and researchers, as cases of cognitive impairment, memory impairment, disorientation and depression have been increasing, evidencing the complexity of the disease^27, 38^.

In this context, here we show that SARS-CoV-2 infects the central nervous system of Syrian hamsters, *Mesocricetus auratus, in vitro* and *in vivo*. More importantly, we observed that the infection actively changes both metabolic and proteomic profile. Proteomic analysis of primary astrocytes in vitro, as well as dissected cortex and hippocampus in vivo, demonstrated relevant changes in proteins related to carbon metabolism, biosynthesis of amino acids and the tricarboxylic acid cycle (TCA). This was further corroborated by the overall increase in glycolysis and oxygen consumption assessed by respirometry, which has already been demonstrated in mononuclear cells and lung epithelial cells. Conversely, KEEG pathway analysis evidenced a conversion to pathways enriched in neurological diseases such as Huntington disease, Parkinson disease and long-term depression, whose symptoms are similar to those observed in many COVID-19 patients.

The fact that viruses actively hijack cellular metabolism is well known, as for CMV, HSV, HIV, vaccinia virus, and many others, as reviewed^39^. This occurs either to facilitate viral replication and virion assembling or to evade local immune responses. Although the presence of replicating SARS-CoV-2 in human brains is still a matter of debate, its impact on neurological function, as memory and cognition, has become unquestionable. In this sense, the relevance of glutamatergic synapses for appropriate brain function is fundamental, as more than 90% of excitatory synapses are mediated by glutamate ionotropic and metabotropic receptors. Due to its importance, glutamate is constantly recycled within the cell, either as a neurotransmitter, as glutamate and aspartate, and transported to pre-synaptic vesicles by vesicular glutamate transporters (Vglut), or following its conversion from glutamine to glutamate and then to alpha-ketoglutarate to fuel the TCA cycle during cataplerosis. This conversion is essential and happens extensively in astrocytes and neurons through the function of glutaminase. Of note, conversion from glucose and glutamine is the major source of glutamate in the brain, as very little is absorbed from the blood circulation. During certain circumstances, this balance may shift from one side to another, such as during infection or neurodegenerative diseases. Thus, it is clear that the balance between glutamate and glutamine plays a pivotal role in both synaptic transmission and energy acquisition for proper neuronal activity, shifting from vesicular neurotransmission to catapletorosis.

Consistently, we showed that SARS-CoV-2 replication in hamster glial cells significantly changes the levels of glutamine and alpha-ketoglutarate, as assessed by metabolomics. Also, glucose and TCA cycle components were reduced, whereas viral replication increases along with cytokine secretion. This is consistent with our previous findings in monocytes^40^. However, here we show that along with increased glycolysis, astrocyte infection with SARS-CoV-2, also used glutamine as a source of carbon. Interestingly, however, viral replication was independent of glycolisys, as 2-DG treatment did not impact viral replication. However, here we show that SARS-CoV-2 replication depends on glutamine, as the blockade of glutaminolysis with 6-Diazo- 5-oxo-L-norleucine (DON) significantly reduced viral replication, and consequently, the production of inflammatory cytokines. This is also consistent with recent reports showing that SARS-CoV2 infection alters the expression of many TCA cycle proteins as well as their metabolites in lung epithelial cells^41^. Thus, it seems clear that SARS-CoV-2 impacts the overall status of protein expression, specifically of those involved in carbon and energy metabolism. And these changes as pivotal for proper brain function.

Among the symptoms observed in patients with long-covid are cognitive impairment, memory decline, and confusion. Conversely, hippocampal glutamatergic neurons as well as a balanced concentration of glutamate or amino acid transporters in the synapses are pivotal for long term potentiation and memory^42^. Thus, it is suitable to speculate that during COVID-19, glutamine consumption due to viral hijacking of cellular metabolism may affect glutamate/glutamine balance, and thus leading to the neurological symptoms mentioned. In fact, it has already been shown that SARS-CoV-2 changes the expression of excitatory amino acid transporters as SLC1A3 in COVID-19 patients astrocytes^43^, which is important for the recapture of synaptic glutamate. Corroborating this hypothesis, a recent case report investigated a 29 years old COVID-19 patient that developed transient attention deficit and memory problems. Conversely, Magenetic Resonance Spectroscopy (MRS) for the analysis of glutamate, glutamine and N-acetyl aspartate clearly showed a reduction of these molecules in the dorso-lateral prefrontal cortex (DLPFC), which is corroborated by our findings. Interestingly, three months after infection, neurological impairment has stope as well as glutamate/glutamine levels were reestablished^44^.

Thus, to our knowledge, this is the first report to demonstrate the impact of COVID-19 over the metabolic and proteomic profile of brain cells, specifically astrocytes. Moreover, we show that SARS-CoV-2 benefits from glutamine as an important source of energy, which in fact has already been shown for vaccinia virus^45^ and cytomegalovirus^46^ in distinct cellular populations. Moreover, our proteomic analysis of hamster’s brains *in vivo* compared to human single cell RNAsequencing corroborates the importance of the metabolic and energetic imbalance of glutamate/glutamine that may take place in the brain of neurological in COVID-19 patients. Our data brings attention not only to the physiology of the COVID-19 brain but also suggesting that glutamate signaling may be target of specific therapies in patients that develop neurological impairments, as memory loss and cognitive decline, as already discussed elsewhere^47^.

---

**Supplementary figure 1:**
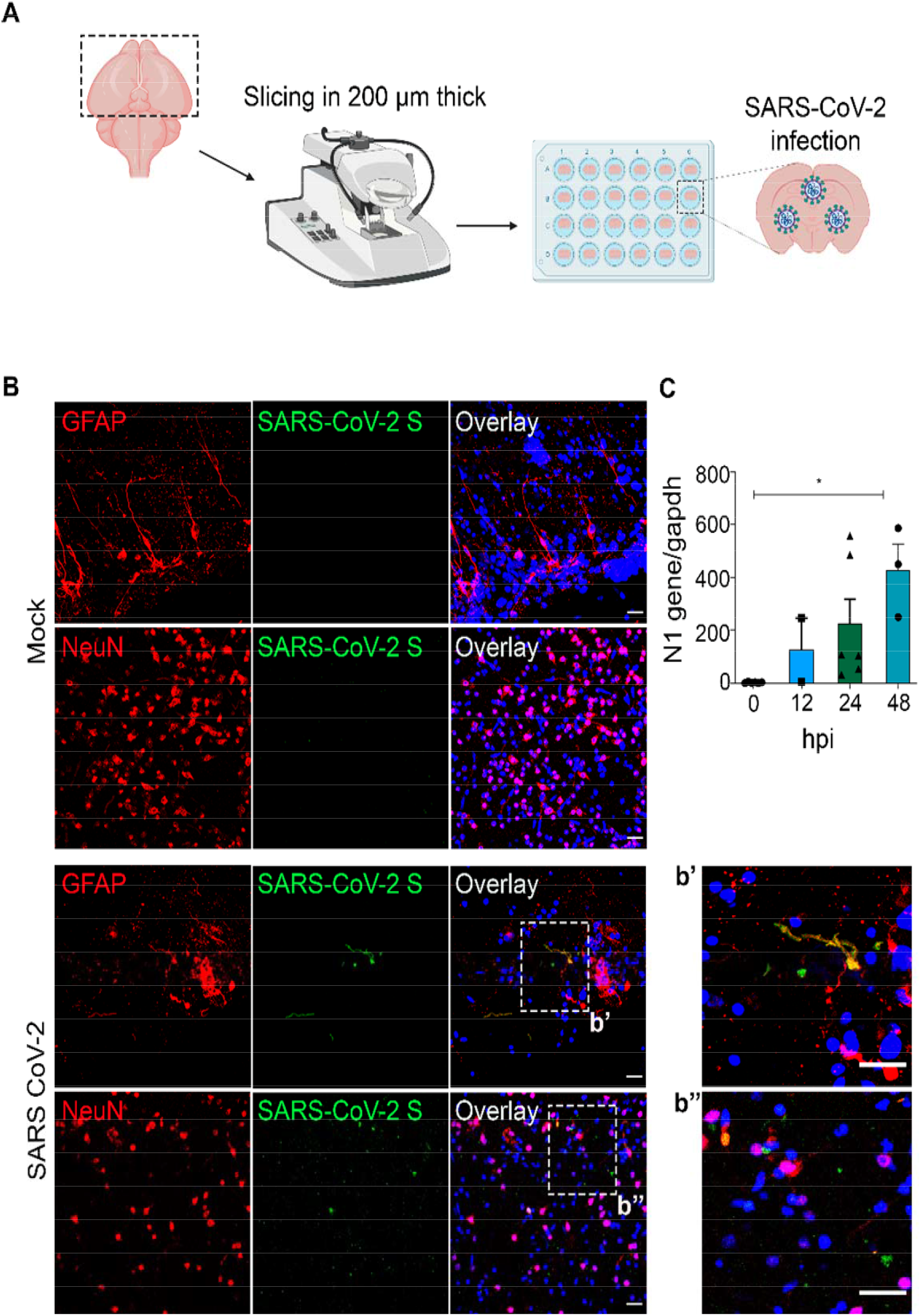
SARS-CoV-2 infection in Syrian hamster brain slices. Scheme illustrating how brain slices were obtained and infected with SARS-CoV-2 3x10^6^ TCID_50_ (**A**) and analyzed for the detection of SARS-CoV-2 spike protein (green) in astrocytes (GFAP) and Neurons (NeuN) (**B**). SARS-CoV-2 nucleocapsid gene was also analyzed at 0, 12, 24 and 48 hpi (**C**). Statistical analyses were performed by *t-test*and *\**p < 0.05 was considered significant.

**Supplementary Figure 2.**
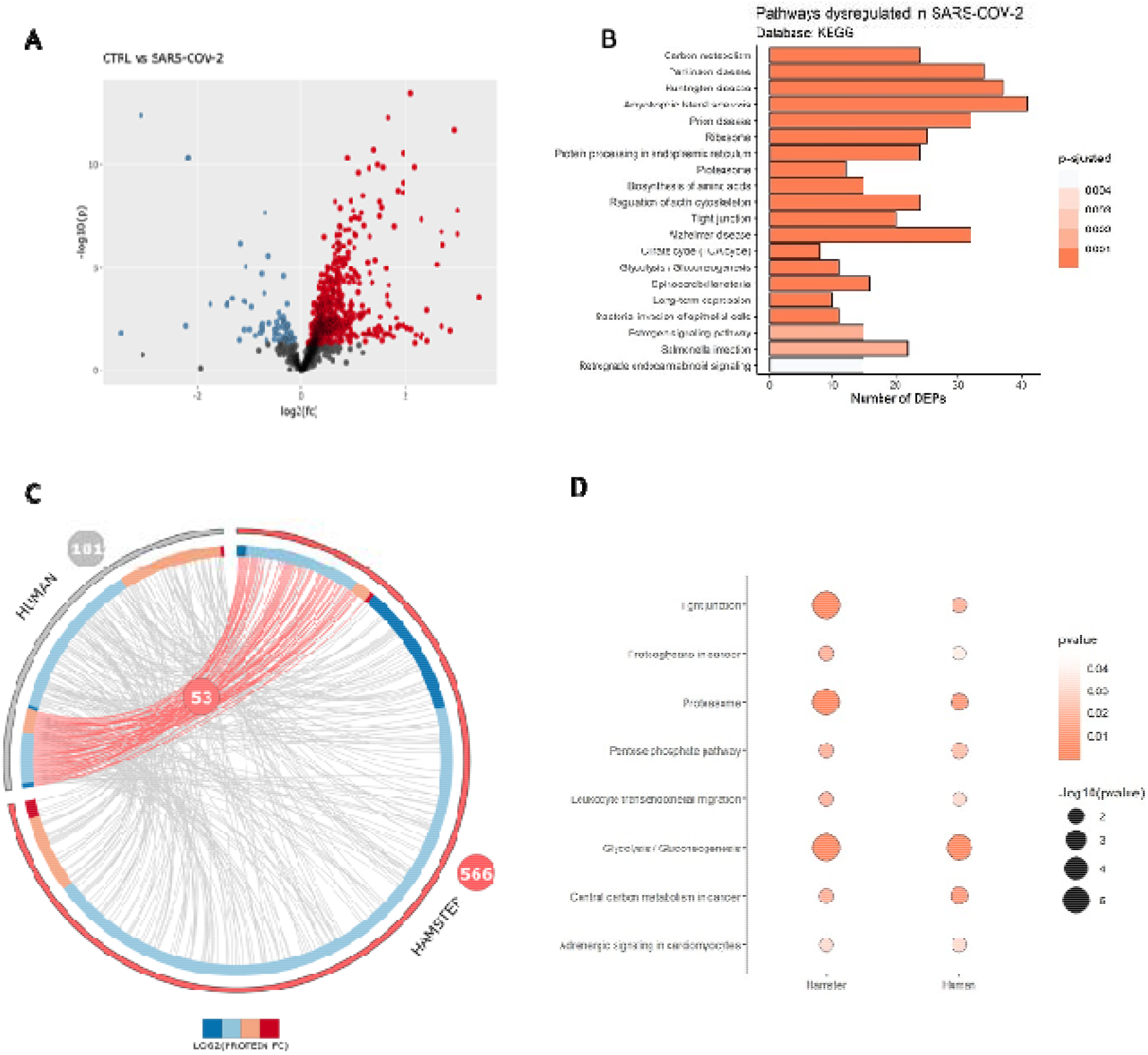
Protein profile hamster’ astrocytes cultures infected with SARS-CoV-2. Volcano plot showing the DEP of mock- and infected groups (**A**). The y axis represents the statistical significance whereas the x axis represents the log2 normalized fold change of the proteins. KEGG pathways analysis of DEPs (**B**). The pathways were ranked by FDR adjusted significance. Circos plot of hamster-to-human primary astrocytes DEP comparison (**C**). The red and gray bars represent up and down regulated proteins, respectively. The red lines connect convergent DEPs between groups. The convergent proteins were fed into a bubble plot (**D**) showing specific enriched pathways and their expression level.

**Supplementary figure 3.**
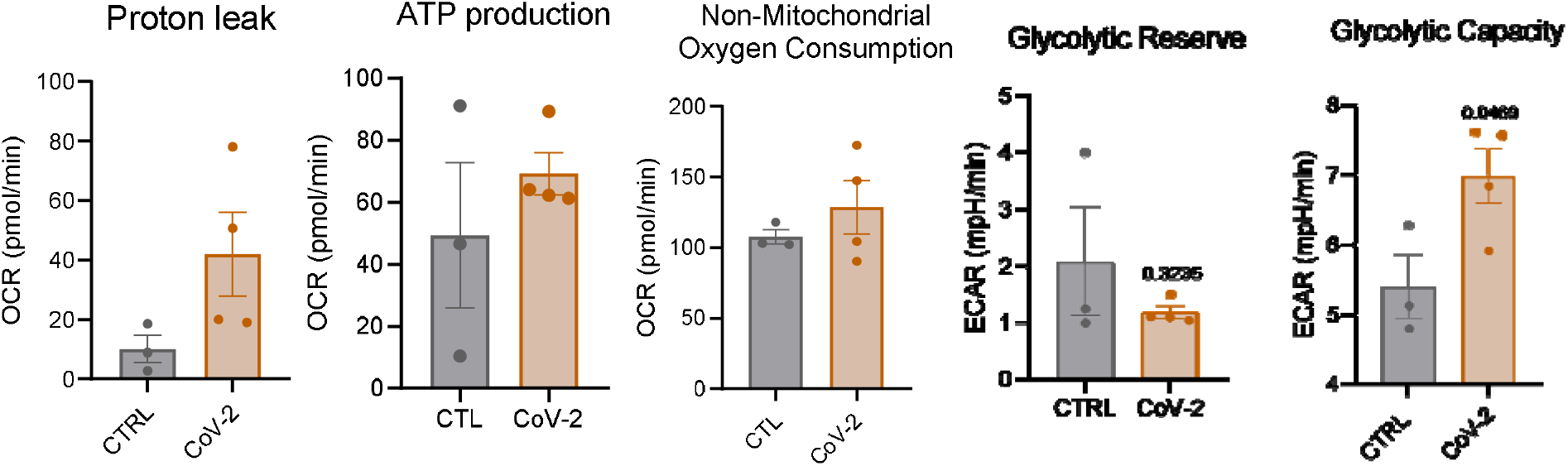
Bioenergetics analysis of SARS-CoV-2 infected hamsters’ astrocytes. Mitochondrial respirometry was performed after SARS-CoV-2 infection (MOI 0.1) at 72 hpi. Bioenergetic profile of oxygen consumption rate (OCR) and extracellular acidification rate (ECAR). Statistical analyses were performed by *t*-test and p < 0.05 considered significant. Cultures were performed in triplicates. Graph represents 2 independent experiments.

**Supplementary figure 4.**
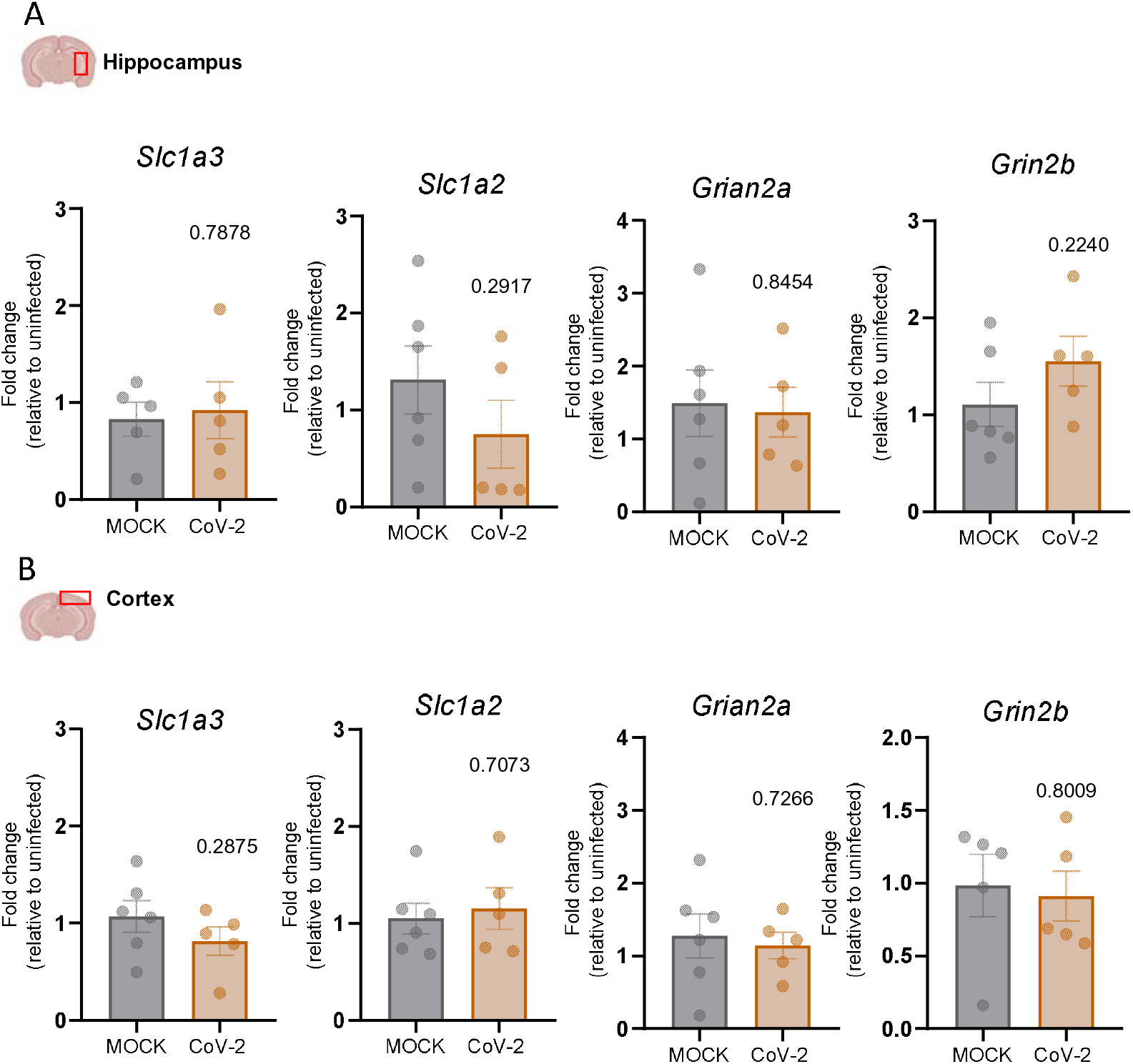
Quantitative gene expression of glutamate transporters and receptors in the brain of SARS-CoV-2 infected hamsters. Quantitative PCR for the expression of amino acid transporters Eaa1 and Eaat2 (*Slc1a3* and *Slc1a2,* respectively) and the subunits of ionotropic glutamate receptor NMDA (*Grian2a* and *Grian2b*). Fold change normalized by the relative expression of uninfected groups. Statistical analyses were performed by *t*-test and p < 0.05 was considered significant.

**Supplementary figure 5.**
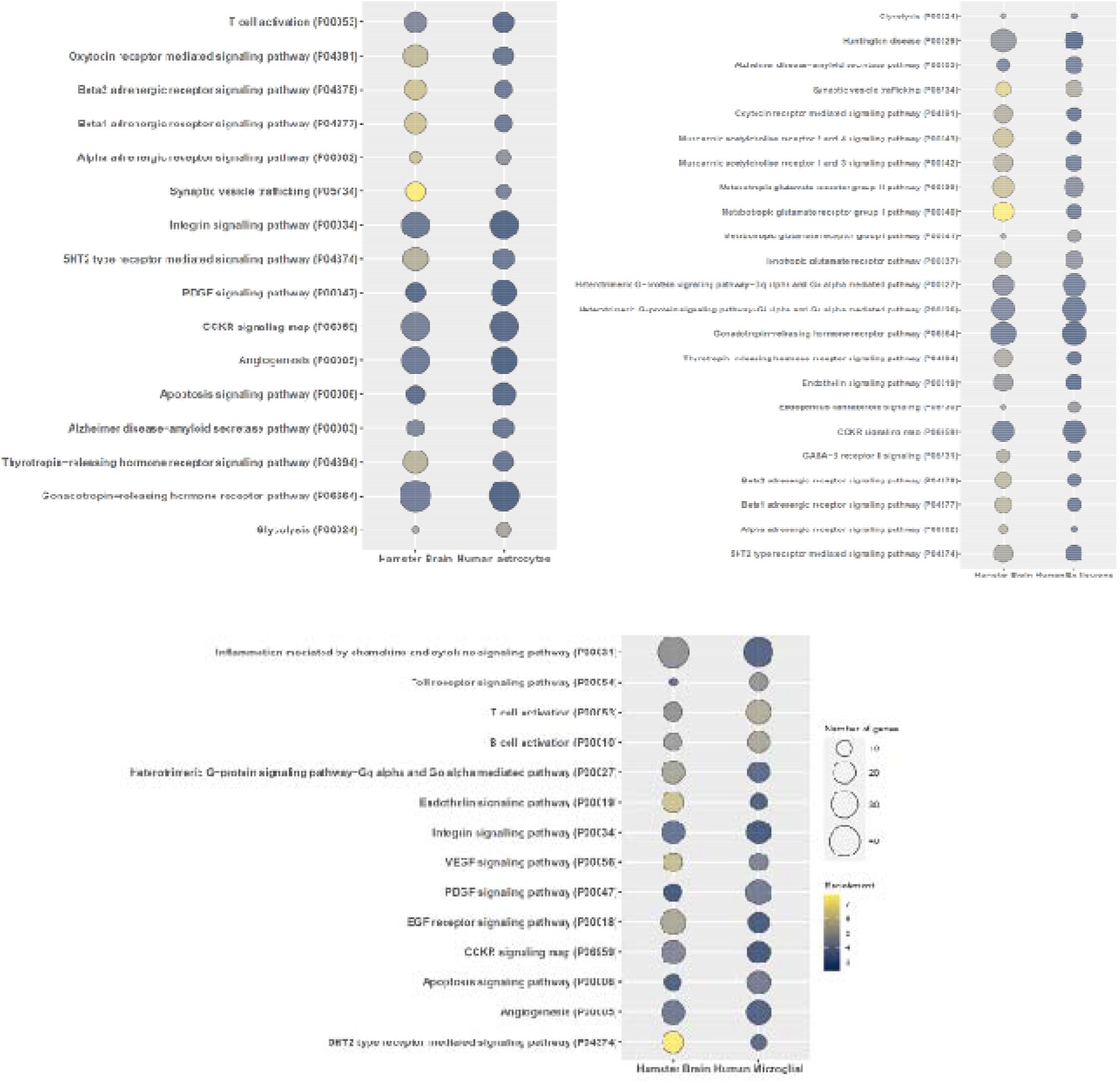
Human and hamster specific brain pathways converge during SARS-CoV-2 infection. Comparisons of the expression and enrichment levels between convergent ontology pathways in hamster proteomic data and human astrocytes (**A**), excitatory neurons (**B**) or microglial cells (**C**) of publicly available single-cell sequencing data. The enrichment level and number of DEPs are represented by size and color of the bubbles.

## Acknowledgements

We would like to thank all the members of the Neuroimmune Interactions Laboratory and the Laboratory of Neuroproteomics for discussions and suggestions. Elia Garcia Caldini from the Laboratory of Cellular Biology and the CNPEN/LNaNo personnel for the support with the TEM images. This research used facilities of the Brazilian Nanotechnology National Laboratory (LNNano), part of the Brazilian Centre for Research in Energy and Materials (CNPEM), a private non-profit organization under the supervision of the Brazilian Ministry for Science, Technology, and Innovations (MCTI). The (names of the facilities) staff is acknowledged for the assistance during the experiments (proposal numbers). No thanks to the federal government, whose huge financial cuts make Brazilian science to agonize.

